# *E. coli* can eat DNA as excellent nitrogen source to grow quickly

**DOI:** 10.1101/2022.03.07.483256

**Authors:** Lili Huang, Yehui Zhang, Xinmei Du, Ran An, Xingguo Liang

## Abstract

Is DNA or RNA a good nutrient? Although scientists have raised this question for dozens of years, few textbooks mention the nutritional role of nucleic acids. Paradoxically, mononucleotides are widely added to infant formula milk and animal feed for improving Intestinal health. Here, found that *E. coli* can grow well in the medium with DNA as carbon and nitrogen sources. More interestingly, in the presence of glucose and DNA, bacteria grew more rapidly, showing that bacteria can use DNA as an excellent nitrogen source. Surprisingly, the amount of DNA in culture media decreased but its length remained unchanged, demonstrating that *E. coli* ingested long DNA directly. The gene expression study shows that *E. coli* mainly ingests DNA before digestion and digests it in the periplasm. *Bifidobacterium bifidum* can also use DNA as the nitrogen source for growth but not efficiently as *E. coli*. This study is of great significance to study DNA metabolism and utilization in organisms. It also lays a foundation to understand the nutritional function of DNA for intestinal flora and human health.

## Introduction

In the natural environment, extracellular or free DNA can be found everywhere, but its utilization by organisms has not been clarified. In the soil, for example, DNA concentration is between 0.08 and 2 μg/g (Nielsen *et al*, 2007); In the ocean, 0.044 mg/L of DNA has been found (Lorenz *et al*, 1994). Extracellular DNA is circulating throughout animal bodies in the blood and other fluids, including urine, milk, amniotic fluid and bronchial lavage (Vlassov *et al*, 2004; Hoskins *et al*, 1978). It is also present at the wound contaminated by bacteria (Brandt *et al*, 1995; Ulmer *et al*, 1996). On the other hand, bacteria are also everywhere and can utilize almost any organic molecules from organisms as “food”. However, few textbooks of nutrition mention the nutritional role of nucleic acids, although scientists have raised the question of “Is DNA or RNA a good nutrient?” for dozens of years. Paradoxically, mononucleotides are added to infant formula milk as the essential component, and RNA-rich formula is widely used in feed of animal offspring. DNA and RNA products have also been used in functional food. One reason for nucleic acids being ignored as the nutrition is that researchers believe its content (especially for DNA) is less than proteins or polysaccharides. Actually, it is a misunderstanding especially in the world of prokaryote. In an *E. coli* cell, for example, nucleic acids account for 7% of the wet weight, and it is only less than protein, which is 15% (Lewin *et al*, 2002). The content of DNA in *E. coli* is about 1.0% (0.5-1.5% wet weight). From the perspective of chemical structure, it is easily to see that nucleic acid and its derivatives are higher in carbon (C), nitrogen (N), and phosphate (P). Sulphur (S) has also been found in some bacteria. For example, Deng et al. have found that the genome of many bacteria involves the sulphur atoms attaching to phosphates on DNA (Zhou *et al*, 2005). Accordingly, it is worth studying nucleic acids deeply from the perspective of nutrition, not just from that of molecular biology and biotechnology. Many exciting findings can be expected in the area of nucleic acid metabolism and nutrition.

Bacteria have been shown to utilize DNA as element sources of P, C and N for growth under special conditions (McDonough *et al*, 2016; Seper *et al*, 2011; Pinchuk *et al*, 2008; Jørgensen *et al*, 1993; Kroer *et al*, *1994;* Paul *et al*, 1988; Redfield *et al*, 1993; Mulcahy *et al*, 2010; Lennon *et al*, *2007*). For example, *V. cholera* has been shown to assimilate phosphate from extracellular DNA and nucleotides, and PhoX, UshA and CpdB have been identified as the major periplasmic phosphatases (McDonough *et al*, 2016); The extracellular deoxyribonucleases of Dns and Xds digest DNA first before assimilation (Seper *et al*, 2011). The genus *Shewanella*, one kind of metal-reducing bacteria, could use extracellular DNA as carbon and energy sources (Pinchuk *et al*, 2008). The possibility of digesting DNA to nucleotides for direct DNA and RNA synthesis was also discussed (Jørgensen *et al*, 1993; Kroer *et al*, 1994; Paul *et al*, *1988;* Redfield *et al*, 1993). Mulcahy et al. showed that *P. aeruginosa* may secrete an extracellular deoxyribonuclease (DNase) to degrade extracellular DNA and utilize it as the C, N and P source (Mulcahy *et al*, 201*0*). Some marine bacteria were also reported to utilize DNA as a nutrient source of C, P and N, but the culture media contained HEPES or other compounds of organic carbon and nitrogen (Lennon *et al*, 2007). *E. coli,* one model bacterium for molecular biology, has also been shown to be capable of consuming DNA as the sole source of carbon and energy (Palchevskiy *et al*, 2006). However, these studies on DNA utilization are scattered and not so persuasive from the viewpoint of nutrition. We did not find any research report using DNA as the nitrogen source in the presence of other carbon source, although the content of nitrogen in DNA is even higher than amino acids. It is well known that glucose is the best carbon source for most organisms, and DNA cannot be a better carbon source than glucose. Actually there are a large number of plants in natural environment to provide potential carbon sources, but the nitrogen source is relatively poor.

From the viewpoint of genomics, bacteria were found to “eat” extracellular dsDNA for recombination (or for “sex”) (Palchevskiy *et al*, 2006; Finkel *et al*, 2001; Dubnau *et al*, 1991; Dubnau *et al*, 1999; Goodgal *et al*, 1982; Kahn *et al*, 1984; Redfield *et al*, 1997). Very interestingly, the competent bacteria can bind and transport long extracellular DNA through their cell envelope and integrate the “eaten” DNA into their chromosome (Palchevskiy *et al*, 2006; Solomon *et al*, 1996; Smith *et al*, 1981). The molecular machines set on the cell surface work for this transportation using the energy from ATP (Finkel *et al*, 2001). More interestingly, a model was proposed that the two DNA strands are separated in the periplasm with one strand broken down there and the other ssDNA strand transferred to the cytoplasm. The phosphatase can remove phosphate groups from 5′- or 3′- nucleotides in the periplasm (McDonough *et al*, 2016; Pinchuk *et al*, 2006; Rittmann *et al*, 2005; Roy *et al*, 1982). Released phosphate can traverse the inner membrane via the phosphate specific transport system (Pst/PhoU) (McDonough *et al*, 2014), and nucleosides can pass through nucleoside transporters (e.g. NupC). There are controversies about the biological significance (for evolution, for repair or for nutrition) of this efficient and active intake of dsDNA (Patrick *et al*, 2013).

In theory, as the most significant biomolecule, DNA should be an excellent nutrient. However, there are not strong evidences, especially considering that the amount of DNA is usually trivial in normal food. To solve this confliction, a breakthrough on understanding DNA nutrition is highly expected. In this study, we found DNA can be utilized as an excellent nitrogen source of *E. coli*, as one of the most important model bacteria. When cultured in the coexistence of glucose and DNA (as the sole nitrogen source), *E. coli* grew unexpectedly quickly. As the nitrogen source, DNA was even comparable to glutamic acid, an amino acid. We also found that DNA was “eaten” directly and efficiently without degradation. We believe that the ability of *E. coli* to assimilate DNA as a nutrient indicates that bacteria utilize DNA very actively as a “delicious” food ingredient of high-quality. A model for how *E. coli* assimilates and utilizes DNA is proposed.

## Results

### *E. coli* uses DNA as the sole carbon and/or nitrogen source

To avoid incorporation or entrainment of other carbon (C) or nitrogen (N) sources, we chose the M9 medium (containing 22.0 mM glucose, 18.6 mM NH_4_Cl, 22.0 mM KH_2_PO_4_, 0.10 mM CaCl_2_, 47 mM Na_2_HPO_4_, 8.5 mM NaCl, 2.0 mM MgSO_4_) as the basal medium, using glucose as the sole C and NH_4_Cl as the sole N source. Glucose is the only organic nutrient in M9 medium. As shown in Fig 1A (red blank triangles), *E. coli* grew well in M9 within the first 12 h (as the logarithmic phase). Very interestingly, when salmon sperm DNA (100-250 bp long) was used to replace NH_4_Cl, *E. coli* grew much more quickly and the logarithmic phase was as short as about 3 h (blue solid triangles, Fig 1A).

**Figure 1.**
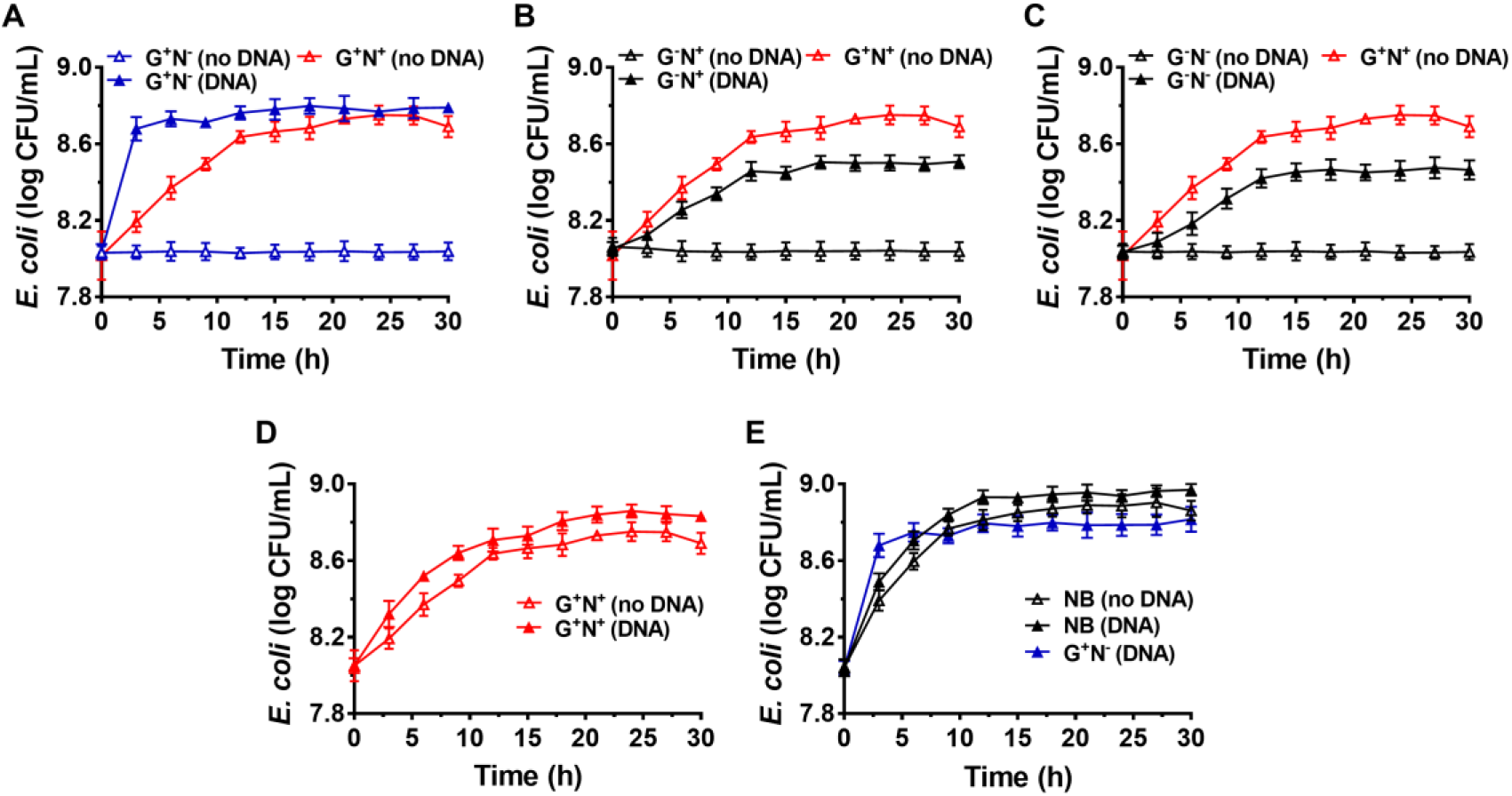
Growth curves for *E. coli* in the presence (DNA) or absence (no DNA) of DNA as the sole carbon and/or nitrogen source in M9 minimal medium. (A) Lacking NH_4_Cl (G^+^N^−^). (B) Lacking glucose (G^−^N^+^). (C) Lacking glucose and NH_4_Cl (G^−^N^−^). (D) Complete M9 minimal medium (G^+^N^+^). For (A-D), M9 medium (G^+^N^+^) was used as the positive control (Red solid triangles). (E) Growth of *E. coli* in nutrient broth (NB) with DNA added as an organic nutrient. For (E), the result is shown for comparison of M9 medium lacking NH_4_Cl (G^+^N^−^) with DNA added as an organic nutrient. The concentrations of DNA and glucose are 0.8 g/L and 4.0 g/L, respectively. Growth of *E. coli* was carried out at 37°C for 30 h. OD_600_ values were measured after certain time intervals. Assays were carried out in triplicate and average values were used (mean and standard deviation are shown).

In the stationary phase, the OD value was even higher than that in the complete medium of M9. Certainly, when neither DNA nor NH_4_Cl was used, no growth of *E. coli* was observed because no N source was present in the medium (blue blank triangles). This effect depends to some extent on the DNA concentration, and 0.2-1.0 g/L DNA shows better effect (Fig EV1B).

In the case that glucose was replaced by DNA (G^−^N^+^, Fig 1B) or both glucose and NH_4_Cl were replaced by DNA (G^−^N^−^, Fig 1C), *E. coli* did not grow so quickly compared with that DNA replacing NH_4_Cl (G^+^N^−^). In addition, the OD values were lower than the positive control (G^+^N^+^, M9 containing glucose and NH_4_Cl) in the stationary phase. Interestingly, even when both NH_4_Cl and DNA were present as the N source, the quick growth as shown in Fig 1A was not observed, and the OD value became a little bit higher than that for M9 (Fig 1D). More interestingly, even for nutrient broth (NB), which is very rich in various nutrients, the growth speed was slower for the first 3 h as compared with the combination of DNA and glucose (see black blank triangles in Fig 1E). Obviously, as compared with NB, M9 medium contains very poor nutrition. Surprisingly, when both DNA and NB were present, the growth speed was also not so quick in the first 3 h (black triangles in Fig 1E). These results show that the extremely quick growth of *E. coli* only occurred in the case that DNA was used as the sole N source (or nutrient poor medium) in the presence of glucose.

It should be noted that, even in the case that DNA was the only organic compound, *E. coli* grew well, although both the growth speed and the final concentration were not so good as that in M9 medium (G^−^N^−^, Fig 1C). This result demonstrates that DNA can be used as the sole carbon and sole nitrogen sources for *E. coli* growth. It is amazing that *E. coli* can grow in the presence of only DNA and inorganic salts. Another exciting result was that even in NB, the addition of DNA could improve both the growth speed and the bacterium density of *E. coli* (comparing black solid triangles and blank triangles in Fig1E), indicating that DNA can be used as a nutrient (probably being mainly used as the nitrogen source even in the case of rich nutrition.

### DNA is comparable to glutamic acid as the nitrogen source for *E. coli* growth

It is well known that amino acids are considered as the best nitrogen source for bacteria among simple organic compounds. Here, we choose glutamic acid for comparison because it is one of the key compounds for metabolism of protein, nucleic acids, and glucose. For more fair comparison, we use 0.8 g/L of DNA (containing about 17% nitrogen in average) or 1.43 g/L of glutamic acid (containing 9.5% nitrogen) with the same concentration of nitrogen (~0.136 g/L). As shown in Fig 2A, when glutamic acid was used to replace DNA, the growth speed (0.30×10^8^ CFU/mL/h) for the first 3 h was not fast as the case for DNA (1.27×10^8^ CFU/mL/h) (comparing triangles and circles). Interestingly, when both glutamic acid and DNA were present, the growth speed became much faster (2.0×10^8^ CFU/mL/h, see squares in Fig 2A). Similar results were obtained when 0.625-5.4 g/L of glutamic acid was used (Fig EV1C). It should be noted that this is different from the cases that DNA was present together with other nitrogen sources such as NH4Cl or components in NB, in which the speed became slower (Fig 1B, 1C, and 1E). These results show again that DNA can be used as an excellent nitrogen source for *E. coli* at least in some special cases.

**Figure 2.**
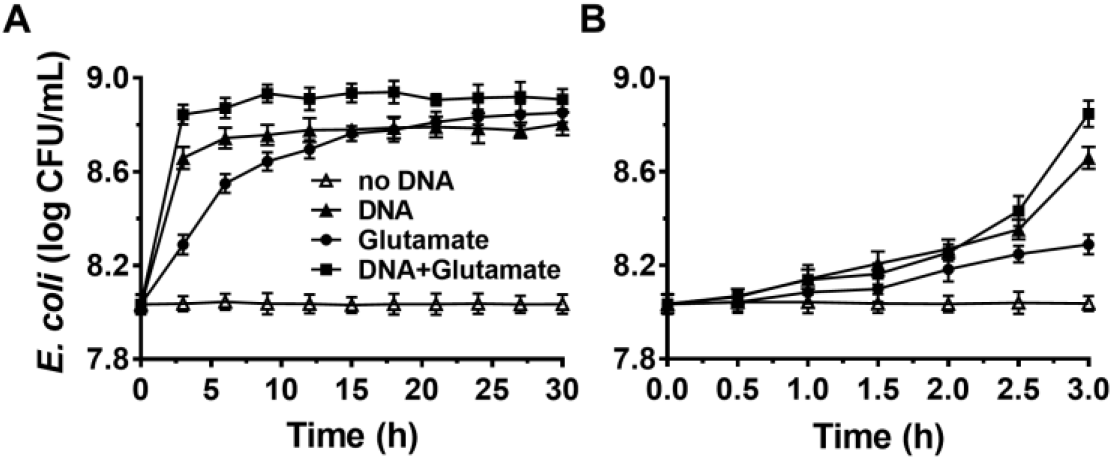
Comparison of DNA with glutamic acid as the nitrogen source for *E. coli* growth in M9 media. (A). The enlarged part of the first 3 h is shown in (B). Solid triangles: 0.8 g/L DNA; Solid circles: 1.43 g/L glutamic acid; Solid squares: both DNA and glutamic acid are present; Blank triangles: no nitrogen source is present. Glucose of 4.0 g/L is used as the carbon source.

As shown in Fig 2B, *E. coli* grew slowly in the first 2 h, indicating that the gene expression of proteins using DNA and/or glutamic acid as nitrogen source was not ready. Obviously, the difference in growth speed was biggest during the 30 min from 2.5 h to 3.0 h of culture. The corresponding growth speeds for glutamic acid, DNA and both glutamic acid and DNA were determined to be 0.46×10^8^, 4.96×10^8^, and 8.80×10^8^ CFU/mL/h, respectively (Fig EV1D). This result shows that there is a synergistic effect for DNA and glutamic acid. In addition, it also indicates that *E. coli* can use DNA as N source very efficiently once *E. coli* prepares well corresponding proteins to digest and utilize DNA. The consumption of DNA and glutamic acid was also estimated. At the stationary phase for DNA as the sole nitrogen source (Fig 2A), about 6.0×10^8^ CFU/mL *E. coli* was obtained. Suppose that the volume of an *E. coli* cell is 0.5 μm^3^ and its dry weight is about 1.5×10^−13^ g, the dry weight of 6.0×10^8^ CFU/mL *E. coli* is equivalent to 0.09 g/L (Fig EV1E). Accordingly, considering the efficiency of anabolism, it can be estimated that 5-20% of organic nutrition (0.8 g/L DNA and 4.0 g/L glucose) was consumed.

### *E. coli* can “eat” DNA directly and grow quickly using DNA as the nitrogen source

How does *E. coli* use DNA in the presence of glucose and grow so quickly? Two possible paths can be proposed. One is that *E. coli* secretes nuclease to digest DNA to oligonucleotides, nucleotides, or nucleosides around the cells, and then these relatively small molecules were ingested. The other one is that *E. coli* ingests long DNA molecules directly without fragmenting them, followed by digestion and utilization in the cells. The latter hypothesis can explain better the quick growth of *E. coli*. To figure out which speculation is correct, we quantitatively analysed the change of DNA amount in the medium at various time intervals.

Fig 3A shows the result of electrophoresis analysis of DNA in the culture media (G+N-, glucose as C source and DNA as N source) after removing *E. coli* cultured for a certain time. It can be seen that the amount of DNA (100-250 bp) decreased to some extent during culture, but the DNA length did not change. For the first 6 h, the intensity of DNA band decreased about 20%, and almost no obvious decrease could be observed after that. Considering that the error for quantitative analysis by electrophoresis is large, we further checked the absorbance (Abs) change of supernatant at 260 nm with the culturing time after removing the *E. coli* by centrifugation (Fig 3B). Interestingly, the Abs change (percentage of Abs decrease based on the Abs at the beginning of culture) increased with a similar trend during culture as that of *E. coli* growth curve (Fig 1A), demonstrating that *E. coli* utilized DNA to grow. Again, Abs did not change much after 6 h of culture, indicating that the utilization stopped or reached the equilibrium between intake and release of molecules with Abs at 260 nm. Decrease of DNA was also observed for other culture media (G^−^N^−^, G^−^N^+^, G^+^N^+^, and NB), and the DNA was not digested completely in all cases (Fig EV2A-D).

**Figure 3.**
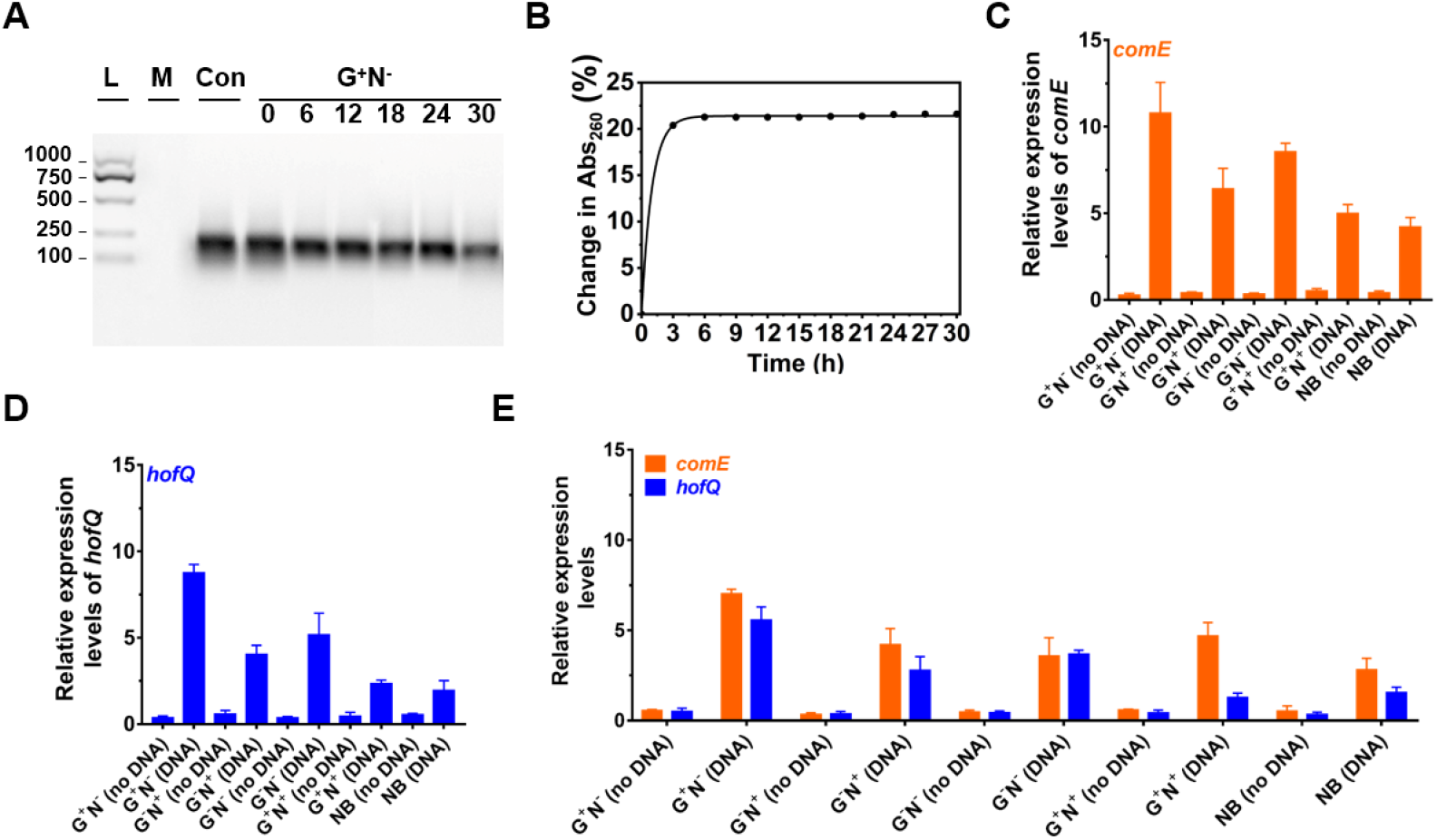
Analysis of DNA uptake to *E. coli*. (A) Analysis of DNA size change by electrophoresis analysis (agarose) at various time intervals of culture in the media using DNA as the sole nitrogen source (G^+^N^−^). L: 100-1000 bp ladder; M: growth media only, Con: control (medium containing DNA without *E. coli*). (B) Change of absorbance at 260 nm at various time intervals of culture (G^+^N^−^). For (A) and (B), the supernatant after centrifugation was used. (C) Expression of *comE* (binding protein for extracellular DNA uptake) in *E. coli*. (D) Expression of *hofQ* (a protein for extracellular DNA uptake) in *E. coli*. For (C) and (D), the *E. coli* after cultured for 3 h was used. (E) Expression of *comE* and *hofQ* genes after 24 h of culture of in various media. *16S rRNA* was used for normalization of gene expression; the relative level was determined using the 2^−ΔΔCt^ method (Landry *et al*, 2014).

We further quantitatively analysed the expressed mRNA of two proteins (ComE and HofQ) related to DNA uptake by reverse transcription quantitative real-time PCR (RT-qPCR). Protein ComE (coding by *comE* gene) is in charge of the DNA binding, and protein HofQ (coding by *hofQ* gene) ingests long dsDNA directly into *E. coli* (Palchevskiy *et al*, 2006; Goodgal *et al*, 1982). As shown in Fig 3C and Fig 3D, the transcription level for both proteins became much higher in the presence of DNA (cultured for 3 h). In the absence of DNA, even for NB medium, the transcription level was also much lower. Interestingly, these two proteins were expressed with the similar order in the presence of DNA: G^+^N^−^ > G^−^N^−^ > G^−^N^+^ > G^+^N^+^ ≈ NB. Obviously, the expression of *comE* and *hofQ* was at a higher level when DNA was used as the sole nitrogen source. Surprisingly, their expression was also up-regulated even when other nitrogen source (NB or NH_4_Cl) was present, indicating that DNA can be ingested once relatively high concentration of DNA (e.g. ≥ 0.8 g/L) is present. It should be noted that DNA accounts for about 0.5% in *E. coli* cells (5.0 g/L).

After 24 h of culture (Fig 3E), even when the growth stopped for G^+^N^−^ (DNA), a high level of expression of *comE* and *hofQ* was kept, although it is a little bit lower than that at 3 h (comparing the data for G^+^N^−^ (DNA) in Fig 3E with those in Fig 3C and 3D). For all the other media containing DNA, the gene expression of *comE* and *hofQ* was also kept at a relative high level (Fig 3E). These results indicate that the presence of high concentration of DNA can stimulate the expression of genes in *E. coli* for DNA uptake. It should be noted that these two proteins only “eat” relatively long dsDNA. Accordingly, it can be concluded that *E. coli* can detect the exogenous DNA and be able to “eat” it as food for growth.

To check whether *E. coli* secretes deoxyribonuclease (DNase) to digest DNA, the expression of *endA* gene (to produce EndA protein, a endodeoxyribonulclease) was investigated. EndA has been proposed to be related to utilizing DNA as nutrition (*Heun et al*, 2012). By gene analysis, we found that the *endA* gene had a high degree of sequence identity with known DNase: *P. fluorescens* extracellular deoxyribonuclease (VVP09331.1). The deduced amino sequence reveals that this DNase has a potential signal peptide (SignalP 5.0 software, score 0.998), demonstrating that it can be secreted outside the *E. coli* cell (Cherny *et al*, 2019). As shown in Fig 4A, when DNA was present in the medium, *endA* was expressed at a much higher level, especially after 3 h of culture. After 24 h, however, the expession level decreased greatly. Certainly, in the absence of DNA in the medium, *endA* was only expressed at a background level, which was even lower than that of *16S rRNA* (as the base for comparison). These results show that the presence of exogenous DNA did improve the expression level of *endA* to digest DNA (see also Fig EV2E-H for the results in other media). However, a new question has arisen that the DNA is digested completely outside the *E. coli*, or in its periplasm. We further tried to clarify this as follows.

**Figure 4.**
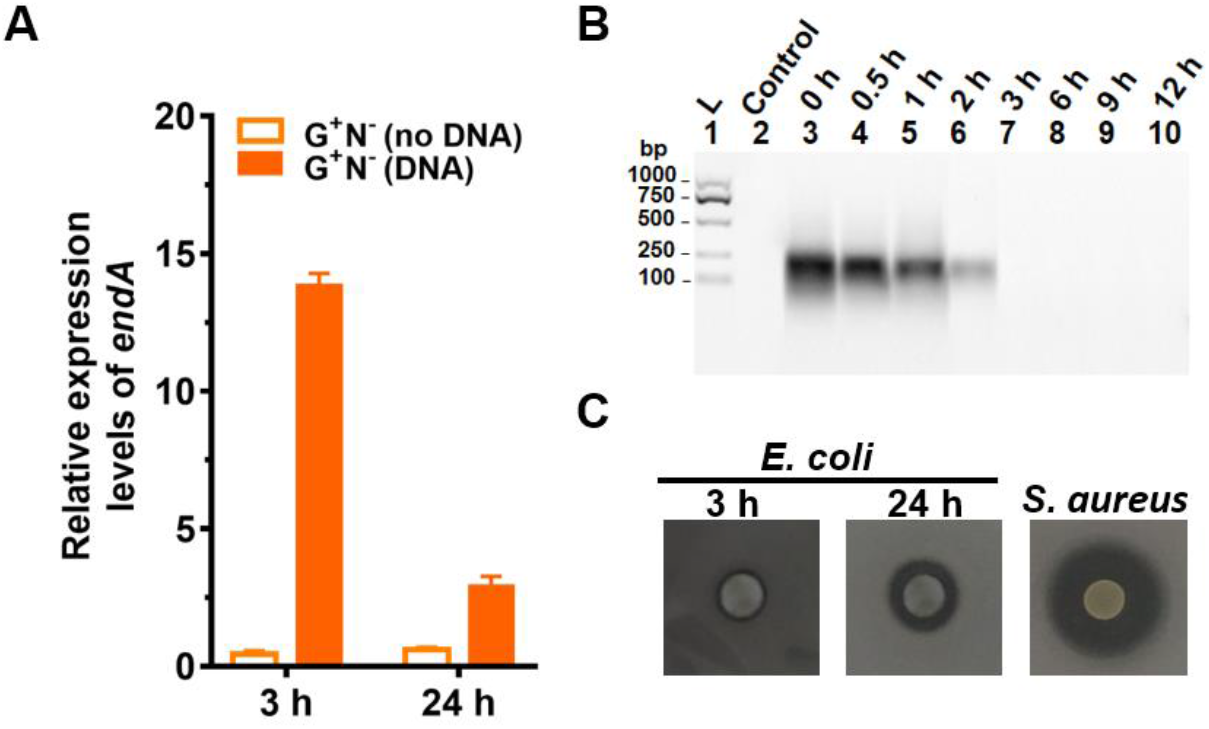
Gene expression and digestion activity of DNase (EndA) from *E. coli*. (A) Gene expression of deoxyribonuclease EndA of *E. coli* in the presence or absence of DNA in the medium (G^+^N^−^).*16S rRNA* was used for normalization, and the relative level was determined using the 2^−ΔΔCt^ method (Landry *et al*, 2014). (B) The digestion of DNA (0.8 g/L) by DNase (0.01 ng/L) at 37°C for various time. (C) Investigation of the extrogenous DNase activity of *E. coli* on agar nutrient broth containing 0.2% (w/v) filter-sterilized DNA. The pictures were taken after incubation at 37°C for 3 h or 24 h. The clear region (black color region here) appeared around the colony reflects the areas where DNA in the agar was digested. *S. aureus* (ATCC6538) for 24 h was used as the positive control.

It is well known that tiny amout of DNase can destory long DNA efficiently, because even 1% digestion occurs, most of 100 bp DNA should be destoryed to shorter ones. We checked this by adding only 0.01 ng/L of DNase I to the medium containing DNA but without *E. coli* (Fig 4B). As expected, no DNA could be observed after 3 h, showing that all DNA was digested to shorter ones. Obviously, the digestion here was much faster as compared with those shown in Fig 3A, indicating that EndA may be mainly present in periplasm but not completely outside the cell. This was further proved by checking the ability of *E. coli* to secrete DNase (EndA) by another approach, in which *E. coli* were spotted onto nutrient broth agar containing 0.2% (2.0 g/L) filter-sterilized DNA and kept for 3 or 24 h (Fig 4C). For the *E. coli* cultured only 3 h, almost no DNase activity was observed, even after 24 h of culutre, the DNase activity was much less compared with the positive control of *S. aureus*.

### Deoxyribonucleotides and deoxyribonucleosides are used as the excellent nitrogen source for E. coli growth

It is easy to imagine that DNA has to be hydrolysed to small molecules (monomers and even smaller ones) for synthesizing macromolecules required for *E. coli* growth. To check whether *E. coli* can ingest directly deoxyribonucleotides (dNMPs) and deoxyribonucleosides, we used dNMPs and deoxyribonucleosides to replace DNA in the medium of G^+^N^−^. As shown in Fig 5A, *E. coli* could grow well using dNMPs in M9 medium without other nitrogen source. Similar as the case using DNA as the sole nitrogen source, *E. coli* grew quickly in the first 3 h. No obvious differences were observed for dAMP, dCMP, dGMP, and dTMP. In the case that deoxyribonucleosides (without phosphate) was used, *E. coli* also grew quickly in the first 3 h. However, the final concentrations of *E. coli* were about 40% lower. For *E. coli* growth, *E. coli* can use the phosphate in dNMPs as P source, but for deoxyribonucleosides media, *E. coli* can only use K_2_HPO_4_ as the P source. Again, no obvious differences were observed for four deoxyribonucleosides (dA, dG, dC, and dT). Similar results were also obtained for other media (G^+^N^−^, G^−^N^−^, G^+^N^+^, NB, Fig EV3)

**Figure 5.**
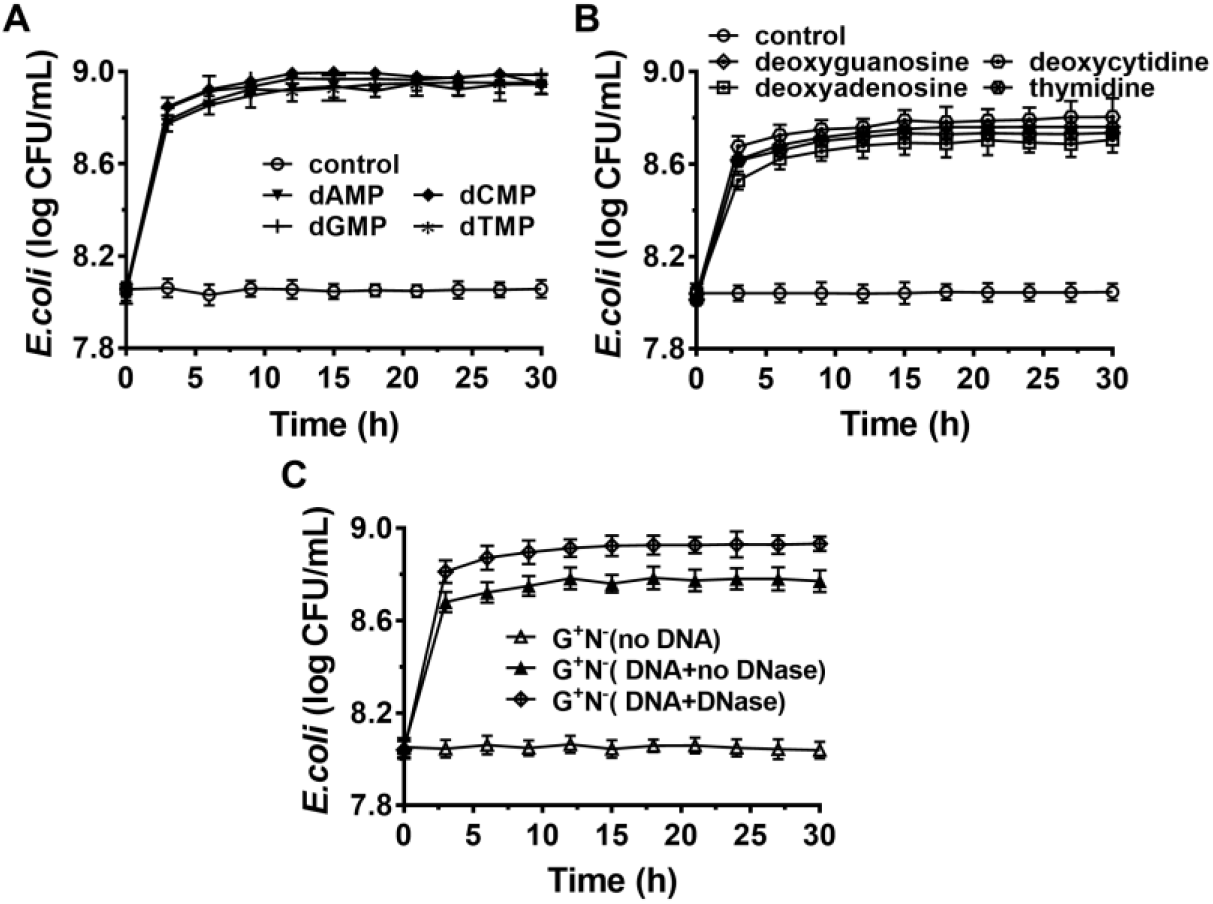
*E. coli* can use deoxyribonucleotides (A) and deoxyribonucleosides (B) as the sole nitrogen source and 4.0 g/L of glucose as the carbon source for growth. The concentration for each deoxyribonucleotide or deoxyribonucleoside was 1.0 g/L. (C) Effect of adding DNase I (0.01 ng/L) to the G^+^N^−^ medium containing DNA on *E. coli* growth.

When 0.01 ng/L of DNase I was added to the cluture medium of G^+^N^−^ (DNA), both the growth speed and the final concentration of cultured *E. coli* increased to some extent (Fig 5C). This result indicates again that *E. coli* can ingest both big dsDNA molecules (> 100 bp) and smaller ones such as dNMPs and deoxyribonucleosides. Similar results were obtained for other media (G^+^N^−^, G^−^N^−^, G^+^N^+^, NB, Fig SEV3). As described previously, in summary, *E. coli* prefers to ingest dsDNA directly and digest it in its periplasm. On the other hand, *E. coli* can also ingest and utilize directly the fragments and monomers of DNA, which are obtained by other factors. These results indicate again that DNA and its digested products of small molecules are all delicious food for *E. coli*.

### *B. bifidum* (gram-positive bacteria) can also use DNA as the sole carbon and/or nitrogen source

At last, we checked whether *B. bifidum* (a gram-positive bacterium) can use DNA as the nutrient (Fig 6). M9 (G^+^N^+^) was also used as the base medium, and NH_4_Cl and/or glucose were removed in some cases (G^+^N^−^, G^−^N^+^, G^−^N^−^). For all cases, *B. bifidum* grew to a stationary stage after 15 to 18 h (Fig 6A, 6B, and 6C). Interestingly, the final concentration of *B. bifidum* in the case that DNA was used as the sole nitrogen source (G^+^N^−^) were higher than other two cases (G^−^N^+^, G^−^N^−^). In G^+^N^−^ medium, although *B. bifidum* grew faster in the first several hours to some extent compared with G^−^N^+^ and G^−^N^−^ media, the very quick growth (in the case of *E. coli*) was not observed. This result indicates that the direct ingestion of dsDNA may be only occurred for *E. coli*, which is a kind of gram-negative bacteria with the periplasmic space. When DNA was additionally added to M9 (G^+^N^+^) or TYP media (medium), no obvious synergistic effect of glucose as carbon and DNA as nitrogen source was observed, indicating again that *E. coli* and *B. bifidum* use DNA in different way. In spite of above results, we can conclude that *B. bifidum* can use DNA for growth, especially in the absence of other nitrogen source.

**Figure 6.**
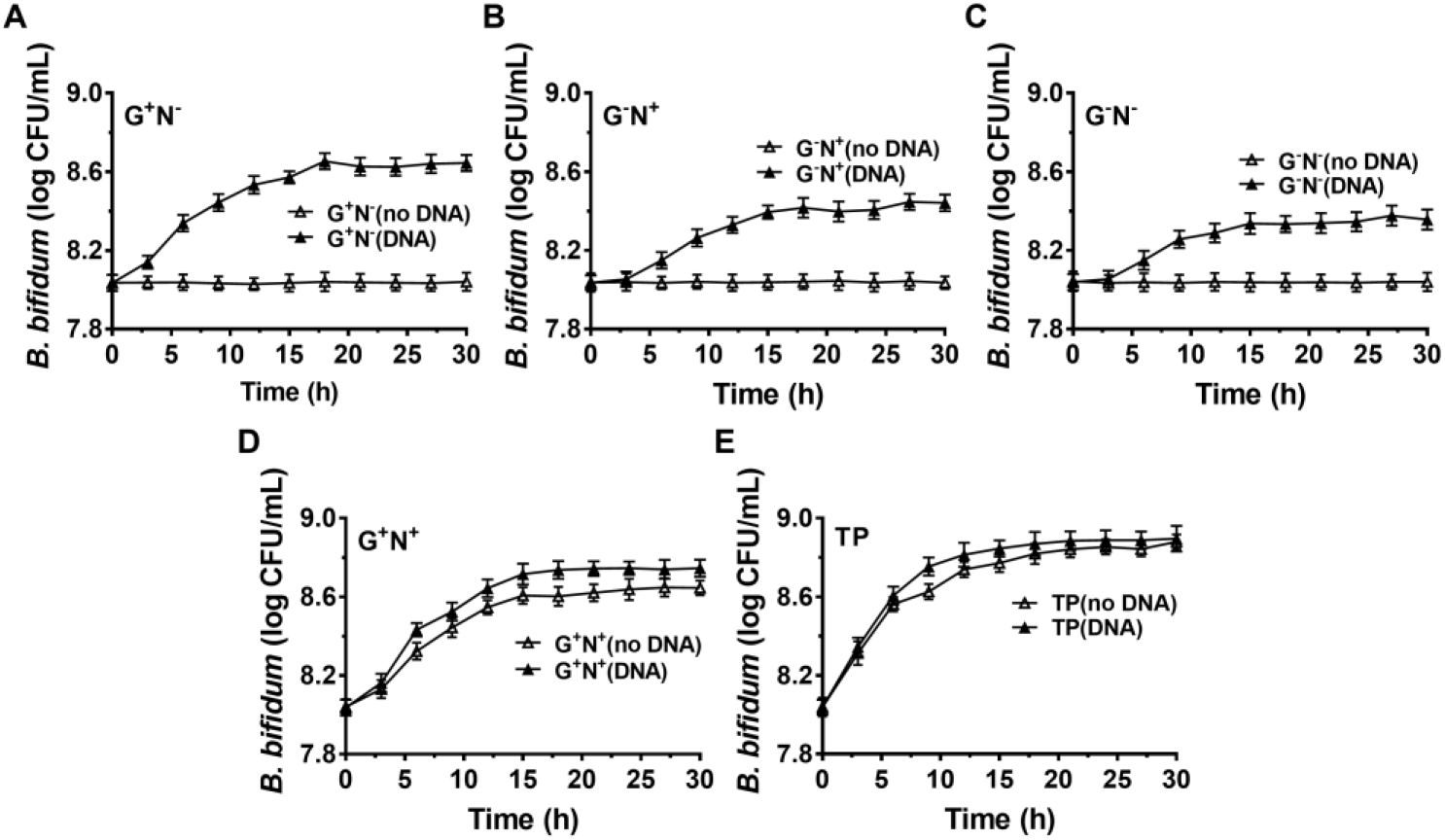
Growth of *B. bifidum* using DNA as the sole carbon and/or nitrogen source. (A, B, C) M9 minimal medium lack of NH_4_Cl (G^+^N^−^), glucose (G^−^N^+^), both glucose and NH_4_Cl (G^−^N^−^), respectively. (D) DNA was added into the full M9 minimal medium (G^+^N^+^). (E) DNA was added in to TPY broth. Concentration of DNA was 0.5 g/L. Media without DNA were used as controls. *B. bifidum* was cultured at 37°C for 30 h under anaerobic conditions. Assays were carried out in triplicate and OD600 values were measured.

## Discussion

Our results clearly show that *E. coli* can ingest dsDNA directly and utilize it as a nutrient, especially as the excellent nitrogen source, which is even comparable to amino acids such as glutamic acid (Fig 1-3). Considering that DNA usually exists in the form of dsDNA either inside or outside a cell (released from organisms), it is more efficient for *E. coli* to “eat” dsDNA to the periplasm and digest DNA there for further utilization. It is less efficient that *E. coli* secretes DNase to the culture medium (outside the cell to the environment) to digest DNA to monomers before ingestion. In addition, secretion of DNase to the environment is not profitable, because *E. coli* usually lives together with many other bacteria. The digested DNA monomers and fragments are easily snatched by *E. coli*’s competitors. On the other hand, the DNase concentration will become much lower if secreted to the culture medium.

Not like proteins or ribose RNA, dsDNA has very simple secondary structure (duplex) so that bacteria are easy to evolve proteins to ingest dsDNA directly. Although *E. coli* can also ingest and utilize directly dNMPs or deoxyribonucleosides as the nitrogen source (Fig 5), it does not mean *E. coli* use this approach (digest dsDNA to small molecules and ingest) as the main one for DNA utilization. On the other hand, it demonstrates that DNA is a good nutrient and *E. coli* may use any kind of DNA and its derivatives. It can be speculated that DNA and its derivatives are preferred to be utilized even when their concentration is much lower than that in the medium we used.

Although many reports claim that bacteria use their own extracellular nucleases to degrade DNA (Seper *et al*, 2011; Pinchuk *et al*, 2008; Mulcahy *et al*, 2010; Heun *et al*, 2012; Cherny *et al*, 2019; Jaskólska *et al*, 2018; Croft *et al*, 1968; Focareta *et al*, 1987; Liechti *et al*, 2013; Alan *et al*, 1978; Gödeke *et al*, 2011; Korczynska *et al*, 2012), it is not clear where these nucleases work outside the cell or in the periplasm. For example, *V. cholerae* has been shown to secrete two nucleases (exonuclease Xds and endonuclease Dns) to break down extracellular DNA into nucleotides as a nutrient (Seper *et al*, 2011; Focareta *et al*, 1987). *Bacillus subtilis* was reported to release DNase to degrade and utilize foreign DNA when it lacks essential nutrients (Alan *et al*, 1978). In this study, we pointed out that nuclease EndA is used as the major nucleases to digest extracellular DNA in the periplasm. EndA (a 235-aa protein) contains an N-terminal signal sequence which is predicted to be located in the periplasm, which is also confirmed experimentally (Heun *et al*, 2012; Cordonnier *et al*, 1965). We believe that for most gram-negative bacteria, their periplasm is the main location for DNA digestion. The weak DNase activity in the media may be caused by the release of DNase from dead bacteria or leak from their outer membrane.

Earlier studies reported the abundance of extracellular DNA in both terrestrial and aquatic environments (Vlassov *et al*, 2007; Finkel *et al*, 2001; Rittmann *et al*, 2005; Trulzsch *et al*, 2001; Dell’Anno *et al*, 1998), where bacteria may use the DNA as a nutrient. Actually, besides those from food, there are two major sources of DNA in the mammalian gut lumen. One is eukaryotic DNA shed from the mucosal epithelium, and the other pool is the DNA from the indigenous microflora. The amount of eukaryotic DNA has been estimated as ~ 5 mg/day in the stomach, 200-500 mg/day in the small intestine, 20-50 mg/day in the colon, and ~6-15 mg/day in the lumen (Croft *et al*, 1968; Croft *et al*, 1973). For some special cases, e.g., acute episodes of diarrhea, eukaryotic DNA can reach 1-10 g/day in the small intestines of humans (Banwell *et al,* 1970; Banwell *et al,* 1971). From the aspect of bacteria, they should not miss these nutrient rich materials (especially rich in nitrogen) as food. In average, the content of nitrogen in nucleic acids is higher than that in amino acids or proteins.

This study has confirmed that salmon sperm DNA can be used as nitrogen and carbon sources by *E. coli*, one of the intestinal bacteria. Especially in the case that no other nitrogen source was present, *E. coli* can utilize dsDNA very quickly in the presence of glucose (Fig 1). Here, very simple media are used because we want to make the results clear without the distribution of other organic nutrients. We believe that, even when *E. coli* lives in a nutrient rich medium, DNA, as well as RNA, can be utilized efficiently. It should be noted that, for most bacteria, DNA and RNA account for 10-20% of their dry weight. Utilization of exogenous nucleic acids to synthesis the DNA and RNA in the bacteria should be energetically favourable. In addition, nitrogen-rich nutrients are usually insufficient compared with carbon source. Accordingly, we believe that the utilization of DNA as well as RNA by intestinal flora is close related to human health. The nucleic acids in the food may be also good nutrient for us.

Based on above analysis, as shown in Fig 7, we proposed a model for the utilization of extracellular DNA by *E. coli*. DNA passes across the outer membrane into the periplasm through porins (HofQ, responsible for transport of DNA across the outer membrane) or nucleoside-specific channel-forming protein (van Alphen *et al*, 1978; Bremer *et al*, 1990). The DNase (EndA) in the periplasm can quickly digest dsDNA to short DNA fragments (<10 bp) or deoxyribonucleotides and deoxyribonucleosides, followed by intake to the cytoplasm of *E. coli*. Usually, a DNA duplex shorter than 10 bp is easily dissociate to ssDNA form (Lehman *et al*, 1962). Accordingly, we believe that the short DNA fragments are mainly transformed to the cytoplasm, but not further digested in the periplasm. The deoxyribonucleotides and deoxyribonucleosides are mainly form in the cytoplasm. Then, these small nitrogen-rich molecules are utilized for synthesis of various molecules for *E. coli* growth by entering the metabolism cycles. The ribose can be used as the carbon source, and nucleobases as the nitrogen source. This speculation is supported by the fact that PDE (phosphodiesterase) has no signal peptide. PDE can only hydrolyse ssDNA in the cytoplasm to yield mono- and dideoxyribonucleotides (Lehman *et al*, 1960). The relatively high level of expression of some corresponding genes to further hydrolyse dNMPs, deoxyribonucleosides, and nucleobases (*xdhA*, *preA*, *pgm* gene) supports the model we proposed (Fig EV4A-E). As expected, expression is also improved for the genes for salvage nucleotide sythesis (*deoD*, *apt, pyrG* gene, Fig EV4F-J) and de novo nucleotide sythesis (*purH, pyrF* gene, Fig EV4K-N).

**Figure 7.**
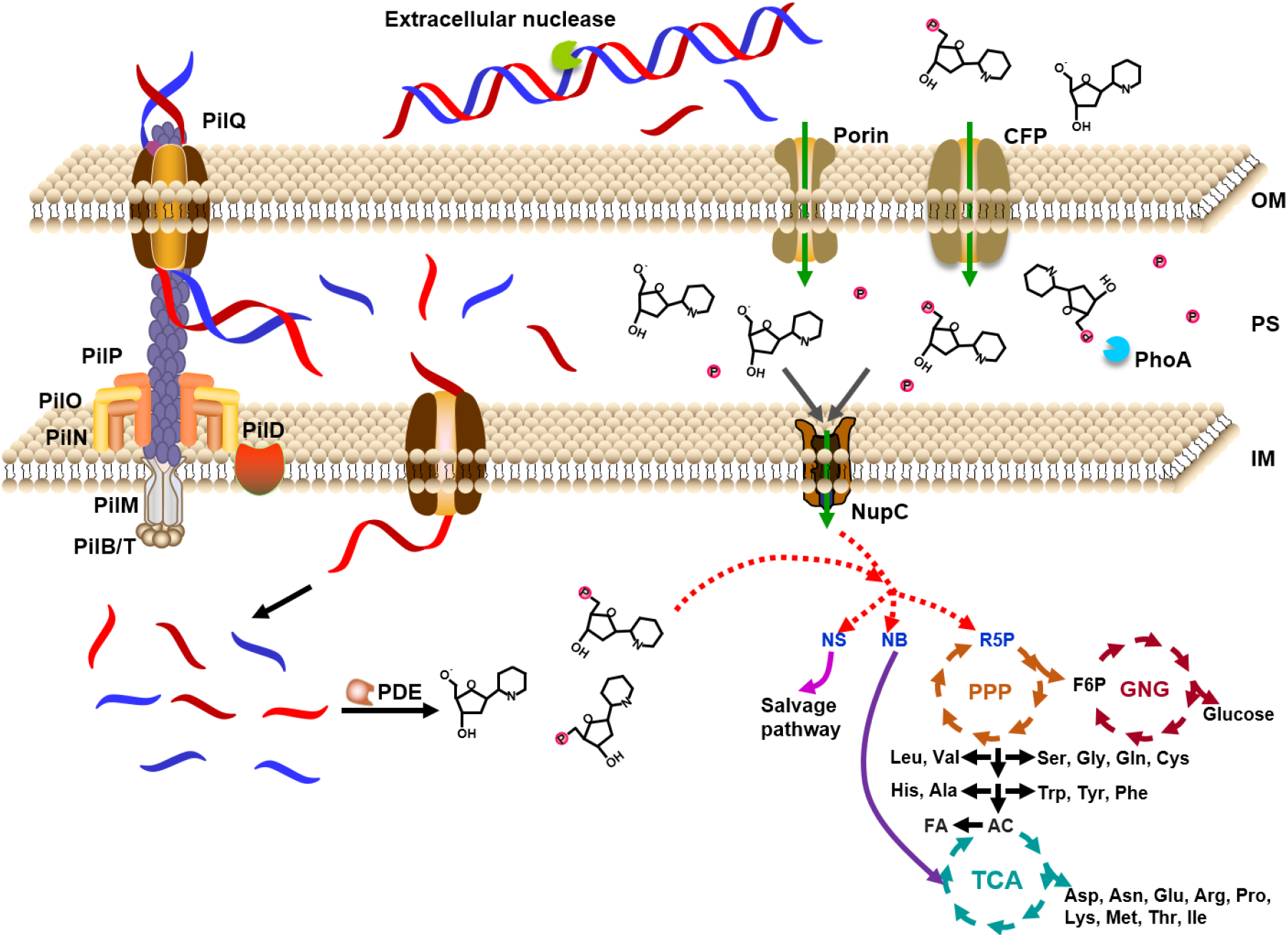
Model for utilization of extracellular DNA as a nutrient by *E. coli*. DNA is caught by proteins on the outer membrane and ingested to the periplasm, where DNase digests DNA to small molecules. The expressed DNase is secreted from the cytoplasm to the periplasm. Small DNA molecules (~7 nt) and monomers are ingested to the cytoplasm by some specific proteins on the inner membrane. After digested to deoxyribonucleosides and even bases and deoxyribose, these small molecules enter the cycles of TCA (Tricarboxylic acid cycle), PPP (Pentose phosphate pathway), and GNG (Gluconeogenesis). The small molecules can also be ingested from the environment directly to the cytoplasm. PDE: *E. coli* phosphodiesterase which hydrolyse deoxyribonucleic acid only in the single-stranded form to yield mono- and dinucleotides; NupC: nucleoside transporter C; CFP: channel-forming protein; NS: nucleoside; NB: nucleobase; R5P: ribose 5-phosphate; F6P: fructose 6-phosphate; AC: acetyl-CoA; FA: fatty acid; OM: outer membrane; PS: periplasm; IM: inner membrane.

The direct ingestion of dsDNA to bacteria has also been reported in a proposed model that the intake DNA is recombined with the bacteria genome (Dubnau *et al*, 1991; Dubnau *et al*, 1999; Goodgal *et al*, 1982; Kahn *et al*, 1984; Solomon *et al*, 1996; Smith *et al*, 1981; Syvanen *et al*, 1998; Stewart *et al*, 1986). In our opinion, this kind of system is mainly used for DNA utilization as a nutrient. Only in the case that the intake dsDNA has almost the same sequences with the bacterium genome, the recombination can occur. It seems that over long evolutionary time, the benefits of maintaining a system for horizontal genetic transfer outweigh the costs. Actually several groups also claim that this dsDNA ingestion system might serve for a nutritional purpose utilizing extracellular DNA (Redfield *et al*, 1993; Redfield *et al*, 1997; Solomon *et al*, 1996; Allemand *et al*, 2012; Burton *et al*, 2010). Therefore, it is quite reasonable that *E. coli* and other organisms take advantage of this system to efficiently “eat” dsDNA.

In conclusion, *E. coli* can utilize DNA as a good nutrient by directly “eat” it. In the periplasm, the eaten DNA is digested and the obtained fragments or monomers are transferred into the cytoplasm as nutrients. The ingested DNA can be used to synthesize almost all the biomolecules for *E. coli*’s reproduction. *B. bifidum* can also use DNA as the nitrogen source and carbon source, although the utilization is carried out in a different way. DNA and its derivatives should be utilized by most organisms as good nutrients. Many scientific new understandings are expected in the area of nucleic acid metabolism and nutrition. Study on utilization of RNA as a nutrient by bacteria is underway in our lab.

## Materials and methods

### Materials

Salmon sperm DNA (GC content is 41.20%, 100-250 bp) was purchased from Tokyo Pharmaceutical Factory Limited (Tokyo, Japan) and solved in water to a concentration of 10 g/L. Purity of nucleic acid was evaluated by UV-Vis spectra. No absorption peak at 280 nm was observed and the Abs_260_/Abs_280_ was higher than 1.8 (Fig EV1A). Oligo DNAs as primers for RT-PCR used in this study (see sequences in Table S1) were synthesized by Sangon Biotech Co., Ltd., (Shanghai, China). *Staphylococcus aureus* (ATCC6538) was purchased from China Center of Industrial Culture Collection (CICC). *E. coli* (No.1.3344) and *Bifidobacterium bifidum* (*B. bifidum*, No.1.5091) were purchased from China General Microbiological Culture Collection Center (CGMCC). Media (Nutrient Broth and TPY broth) were purchased from Hope Bio-technology (Qingdao, China). Nutrient Broth (pH 7.0±0.2) contains the following components (per liter): peptone 10.0 g, beef extract powder 3.0 g, NaCl 5.0 g; TPY broth (pH 6.5±0.1) contains (per liter) casein hydrolysate 10.0 g, peptone from soybean meal 5.0 g, yeast extract 2.0 g, glutose 5.0 g, L-cysteine monohydrochloride 0.5 g, K_2_HPO_4_ 2.0 g, MgCl_2_ 0.5 g, ZnSO_4_ 0.25 g, CaCl_2_ 0.15 g, FeCl_3_ 0.10 mg, Tween-80 1.0 mL. Glutamic acid (98.5%) was form Sinopharm Co., Ltd. (Beijing, China). For all plates, agar was added to a concentration of 15.0 g/L. DNase was purchased from Tiandz Gene Technology Co., Ltd. (Beijing, China). Other chemical reagents such as HClO_4_, phenol/chloroform/isoamyl alcohol were from Solarbio Science & Technology Co., Ltd. (Beijing, China). The RNA prep pure kit and FastKing RT Kit for RNA extraction and reverse transcription were purchased from Tiangen Biothch Co., Ltd. (Beijing, China).

### Culture of bacterial strains in various media

First, *E. coli* were cultured in nutrient broth (NB) at 37°C under atmospheric conditions whereas *B. bifidum* were cultured in TPY (TP) broth at 37°C under anaerobic conditions in anaerobic incubator (Mitsubishi Gas Chemical, Tokyo, Japan). To ensure that no other residual source of carbon and nitrogen remained in the cultures, bacterial strains were then propagated in M9 minimal medium with glucose as the source of carbon and NH_4_Cl as the source of nitrogen. This defined growth medium (pH 7.0±0.2) contains 22.0 mM glucose (4.0 g/L), 18.6 mM NH_4_Cl, 22.0 mM KH_2_PO_4_, 0.10 mM CaCl_2_, 47.0 mM Na_2_HPO_4_, 8.5 mM NaCl, 2.0 mM MgSO_4_.

For nucleic acid-dependent growth using various nucleic acid derivatives (including DNA, deoxyribonucleotides (dNMPs), and deoxyribonucleosides) as the sources of carbon and nitrogen, nucleic acid was added to the carbon or/and nitrogen-free medium to a final concentration ranging from 0.10 to 10.0 g/L. The nucleic acid was added to the medium prior to filtering through a 0.22-μm-pore-size membrane (Jinteng, Tianjin, China). To remove any potential contaminating proteins present in the commercial DNA preparation, DNA was digested by proteinase K (20 g/L) for 30 min, followed by extraction using phenol/chloroform/isoamyl alcohol.

### Growth curves of *E. coli* with DNA, deoxyribonucleotides and deoxyribonucleosides

Initially, the overnight cultures were taken out from the incubator and put on ice to block the growth of bacteria. After cooling on ice for 15 min, the colonized bacteria strains were centrifuged (4500 g, 4°C, 10 min) and washed thrice with 10.0 mL of the corresponding basal medium (without DNA, dNMPs or deoxyribonucleosides). Then the cells were harvested, resuspended in the basal medium, followed by addition of the nucleic acid. All bacterial strains were adjusted by the medium to an optical density (OD) of 0.05 (approximately 1.0 × 10^8^ CFU/mL) at 600 nm, and then incubated aerobically (for *E. coli*) or anaerobically (for B*. bifidum*) at 37°C, respectively. For analysing samples cultured for various time intervals, each of the bacteria suspensions (200 μL) was taken and added into a 96-well plate and measured the OD at 600 nm and 37°C immediately (in some cases if the OD value exceeds 0.8, the samples are further diluted to the OD value between 0.1-0.8). All growth experiments were independently repeated at least thrice. The error for OD values is within 4-18%. Cultures grown in Nutrient Broth (NB), TPY broth or M9 minimal medium lack of DNA, dNMPs or deoxyribonucleosides were served as the control.

### Change in absorbance at 260 nm of exogenous nucleic acid during culture

Cultures were grown in Nutrient Broth, TPY broth and corresponding basal medium (containing nucleic acid) at 37°C for the same cultivation time intervals. The cell-free supernatants (CFS) were obtained by centrifugation of the strains at 13,400 g for 10 min (4°C), followed by filter-sterilization using a sterile filter with the pore-size of 0.22 μm. Absorbance (Abs) at 260 nm was measured using an ultraviolet spectrophotometer (Nanodrop ND 2000, Thermo Scientific). Percentage for Abs change was calculated as [(A_c_-A_t_)/A_c_×100%)], where A_c_ and A_t_ are absorbance values measured at 0 h and after incubation of a certain time. The media containing nucleic acid of the same concentration were used as control (Ac).

### Nucleic acid degradation assays by agarose gel electrophoresis

The CFS of each strain previously filter-sterilized was used to check its activity for nucleic acid digestion. After incubation for a certain time, the DNA left (without digestion) was analysed on a 1.2% agarose gel. The reaction yield was obtained by quantitative analysis of the decrease of the intact DNA band (Molecular Imager Gel Doc XR+ imaging system, Image Lab 3.0). The digestion yield was calculated by comparing the intensity of the band with that before digestion.

### Evaluation of extracellular DNA degradation activity using an agar plate assay

Extracellular DNA degradation activity of *S. aureus* and *E. coli* were examined using a previously described method (31). Briefly, a 10.0 μL aliquot of an overnight culture (washing with PBS buffer (pH 7.2), then resuspend in PBS buffer) was spotted onto nutrient broth agar plates containing 2.0 g/L of DNA, and incubated overnight at 37°C. For visualization, plates were flooded with 3.0 mL of perchloric acid solution (50% v/v perchloric acid in distilled water), and kept for 5 min. DNA-digested areas appeared as a clear zone around the colony against opaque background. The extracellular DNA degradation activity was determined accordingly, because there is a direct relation between the diameter of cleared zone and nuclease activity (Morita *et al*, 2014).

### RNA isolation and reverse transcription PCR

Total RNA was extracted from bacteria using RNA preparation pure kit according to the manufacturer’s instructions. RNA samples were diluted by RNase-Free distilled water, and the optical density (OD) values at 260 nm and 280 nm were used to evaluate the RNA concentration and purity. Next, to determine the mRNA expression of each gene, a total of 10.0 μL reverse transcription system was used as per the instructions of the FastKing RT Kit. The generated cDNA was stored at −20°C. RT-qPCR was performed on a Piko real 96 Real-time PCR System (Thermo Scientific, USA). 16S rRNA was used for normalization of mRNA expression, the relative level of each mRNA was determined using the 2^−ΔΔCt^ method (Landry *et al*, 2014). The PCR conditions were comprised of pre-denaturation at 95°C for 10 min, 40 cycles of denaturation at 95°C for 15 s, annealing at 60°C for 30 s and extension at 72°C for 30 s. All RT-qPCR experiments were conducted with 3 duplicate wells, and each gene experiment was repeated 3 times.

### Statistical analysis

Data are presented as means ± SEM. The GraphPad PRISM 7.0 software (GraphPad Software Corporation, USA) was used for analysis of the experimental data. The analysis of variance (ANOVA) was used to estimate the statistical parameters.

## Data availability

**Expanded View** for this article is available online.

## Acknowledgments

This work was supported by Fundamental Research Funds for Co-construction of Universities in Qingdao to X.L.; Natural Science Foundation of Shandong Province, China ZR2019BC096 to R.A.; and National Natural Science Foundation of China 31571937 to X.L.

## Author contributuions

X.L. and L.H. developed and designed the experiments for the study. L.H. and Y.Z. performed the experiments. L.H., X.D., and X.L. analyzed and interpreted the data. X.L., R.A., and L.H. wrote the paper. All authors reviewed the paper.

