## Supplementary material for "*E. coli* can eat DNA as excellent nitrogen source to grow quickly": Fig EV

*Lili Huang et al*

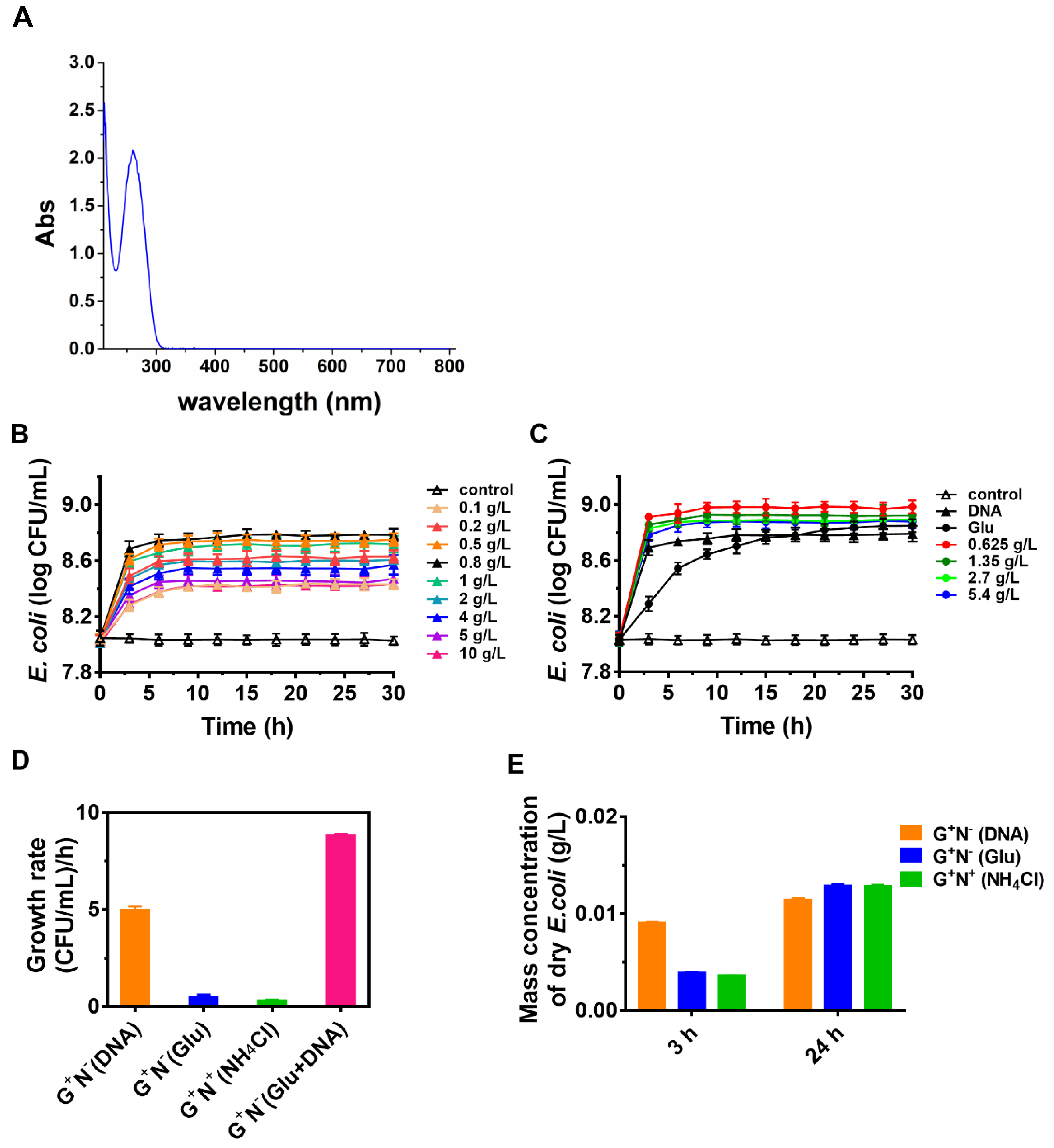

**Figure EV1. Growth of *E. coli* with DNA as the sole nitrogen source.**

(A) Absorbance spectra of salmon sperm DNA used in this study. Nanodrop was used for this analysis. The concentration of DNA is 1.0 g/L. No side peak was observed, showing that the purity is quite high.

(B) Various concentrations of DNA (0.10-10.0 g/L) were used as the sole nitrogen source.

(C) The concentration of DNA was 0.8 g/L, and glutamic acid concentration was 0.625-5.4 g/L. It seems that *E. coli* prefers to use DNA even in the presence of glutamic acid.

(D) Growth rate of *E. coli* with DNA after culture using as the nitrogen source (0.8 g/L) and glucose as carbon source (4 g/L).

(E) Estimated mass concentration of dry *E. coli* after culture using as the nitrogen source (0.8 g/L) and glucose as carbon source (4 g/L). The growth rate of *E. coli* was calculated for the culture from 2.5 h to 3 h in various media. The DNA utilization percentage after culturing at 37°C for 3 h and 24 h was estimated as follows. Assuming the volume of one *E. coli* (containing 70% water) is 0.5×10^-12^ mL (0.7 μm diameter, 1.5 μm length), its dry weight can be calculated as about 1.5×10^-13^ g. The estimated mass concentration (g/L) of dry *E. coli* can be simply calculated by multiplying the number of *E. coli* in one litre.

*Lili Huang et al*

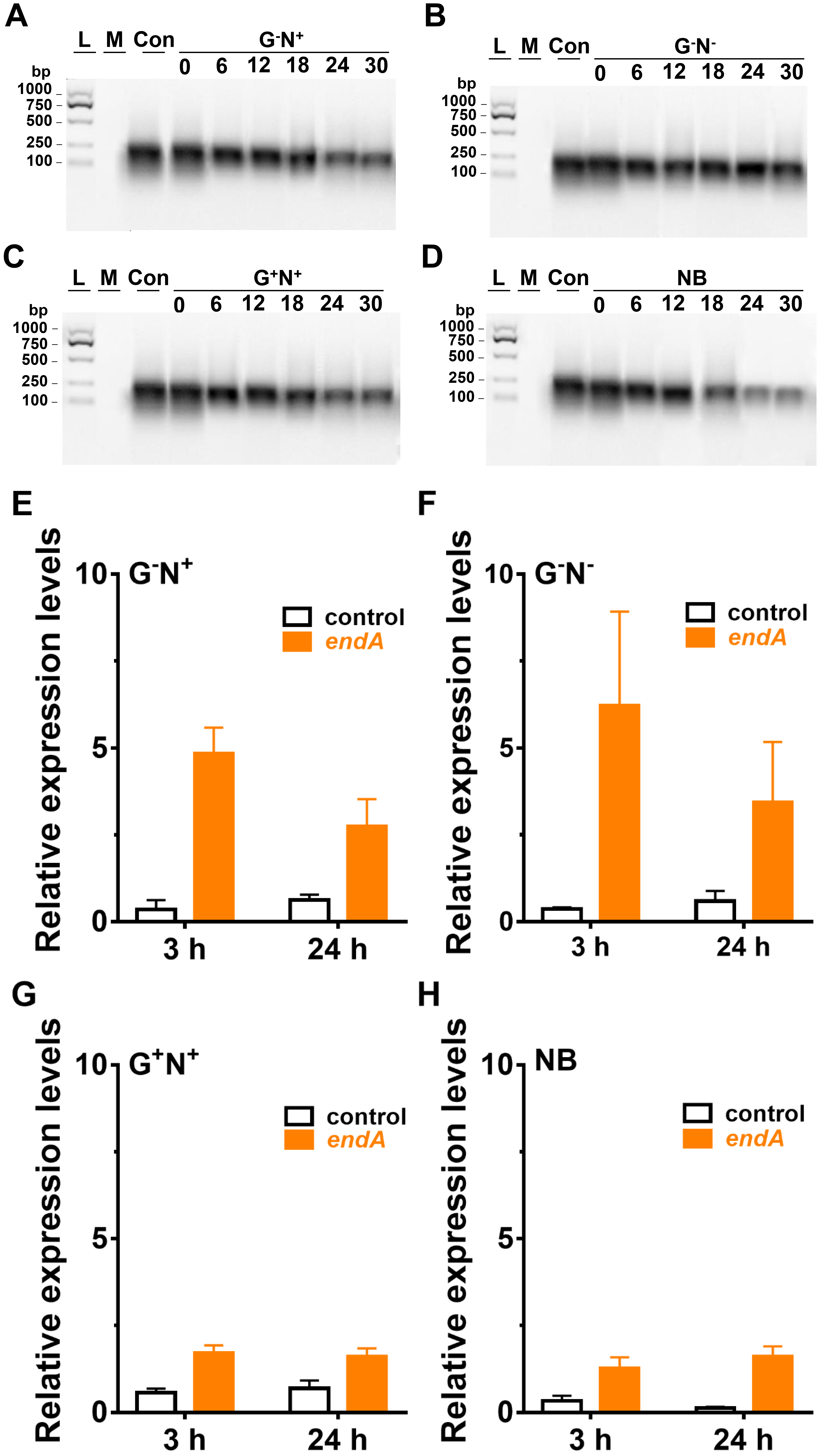

**Figure EV2. Decrease of DNA concentration and gene expression of *endA* of *E. coli* in various M9 media.**

(A) Decrease of DNA concentration in M9 medium lacking glucose (GˉN^+^) after culture of *E. coli*. L: 100-1000 bp ladder; M: growth media without DNA; Con: control (medium containing DNA without *E. coli*).

(B) Decrease of DNA concentration in M9 medium lacking glucose and NH_4_Cl (GˉNˉ) after culture of *E. coli*.

(C) Decrease of DNA concentration in M9 medium (G^+^N^+^) after culture of *E. coli*.

(D) Decrease of DNA concentration in Nutrient broth (NB) instead of M9 after culture of *E. coli*.

(E) Gene expression of *endA* of *E. coli* after clutured in M9 medium lacking glucose (GˉN^+^) containing DNA for 3 h and 24 h.

(F) Gene expression of *endA* of *E. coli* after clutured in M9 medium lacking glucose and NH_4_Cl (G^ˉ^Nˉ) containing DNA for 3 h and 24 h..

(G) Gene expression of *endA* of *E. coli* after clutured in M9 medium (G^+^N^+^) containing DNA for 3 h and 24 h.

(H) Gene expression of *endA* of *E. coli* after clutured in nutrient broth (NB) instead of M9 containing DNA for 3 h and 24 h.

*Lili Huang et al*

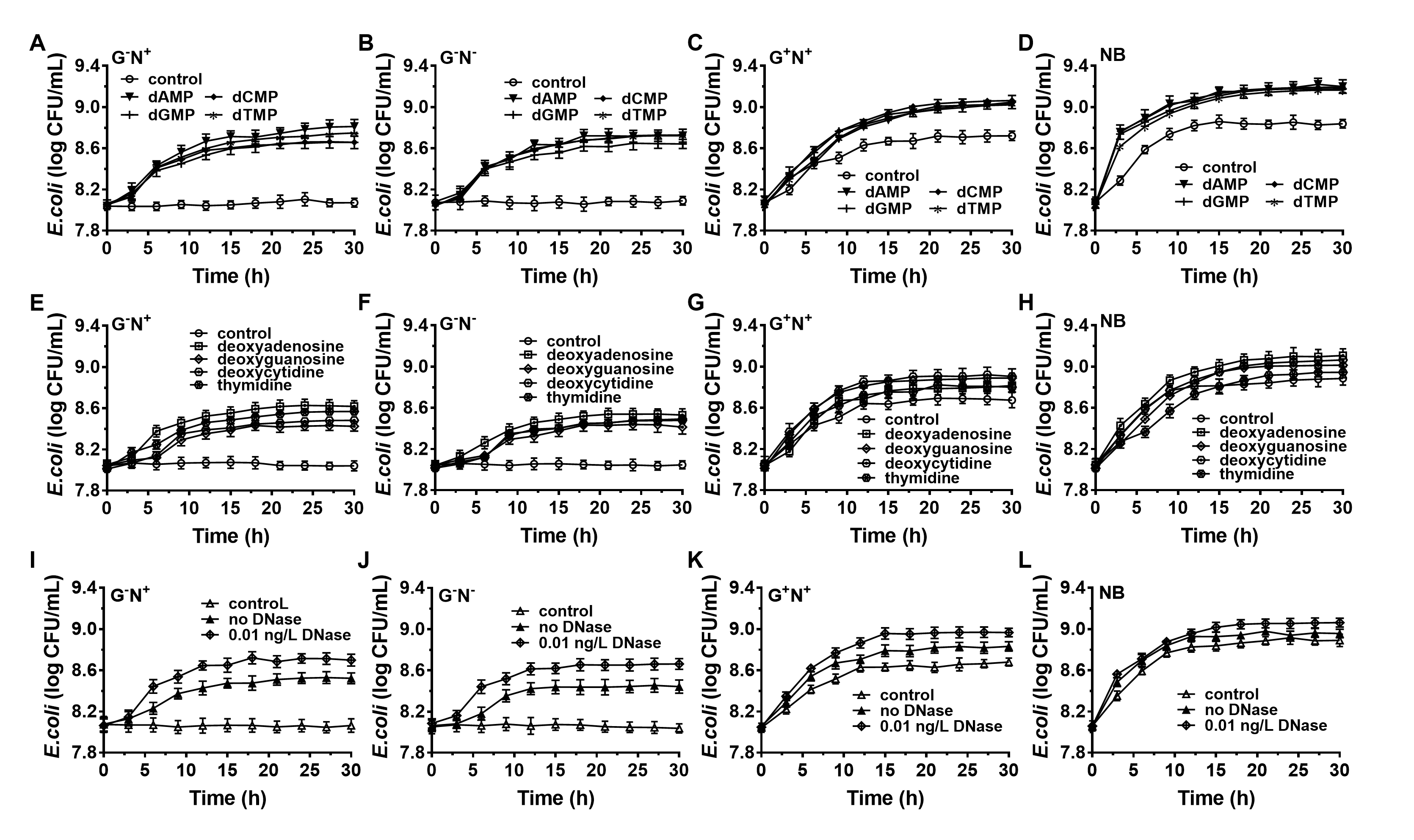

**Figure EV3. Time course of E. coli growth in various M9 media containing nucleic acid derivatives.**

(A)Time course of *E. coli* growth in M9 medium lacking glucose (G^ˉ^N^+^) containing deoxyribonucleotides (1.0 g/L) as the sole carbon and/or nitrogen source.

(B) Time course of *E. coli* growth in M9 medium lacking glucose and NH_4_Cl (G^ˉ^Nˉ) containing deoxyribonucleotides (1.0 g/L) as the sole carbon and/or nitrogen source.

(C) Time course of *E. coli* growth in M9 medium (G^+^N^+^) containing deoxyribonucleotides (1.0 g/L) as the sole carbon and/or nitrogen source.

(D) Time course of *E. coli* growth in nutrient broth (NB) instead of M9 containing deoxyribonucleotides (1.0 g/L) as the sole carbon and/or nitrogen source.

(E) Time course of *E. coli* growth in M9 medium lacking glucose (G^ˉ^N^+^) containing deoxyribonucleosides (1.0 g/L) as the sole carbon and/or nitrogen source.

(F) Time course of *E. coli* growth in M9 medium lacking glucose and NH_4_Cl (G^ˉ^Nˉ) containing deoxyribonucleosides (1.0 g/L) as the sole carbon and/or nitrogen source.

(G) Time course of *E. coli* growth in M9 medium (G^+^N^+^) containing deoxyribonucleosides (1.0 g/L) as the sole carbon and/or nitrogen source.Complete.

(H) Time course of *E. coli* growth in various M9 media lacking glucose and NH_4_Cl (G^ˉ^Nˉ) containing deoxyribonucleosides (1.0 g/L) as the sole carbon and/or nitrogen source.Nutrient broth (NB) instead of M9.

(I) Time course of *E. coli* growth in M9 medium lacking glucose (GˉN^+^)containing DNA in the absence and presence of DNase (0.01 ng/L).

(J) Time course of *E. coli* growth in M9 medium lacking glucose and NH_4_Cl (GˉNˉ) containing DNA in the absence and presence of DNase (0.01 ng/L).

(K) Time course of *E. coli* growth in M9 medium (G^+^N^+^) containing DNA in the absence and presence of DNase (0.01 ng/L).

(L) Time course of *E. coli* growth in nutrient broth (NB) instead of M9containing DNA in the absence and presence of DNase (0.01 ng/L).

*Lili Huang et al*

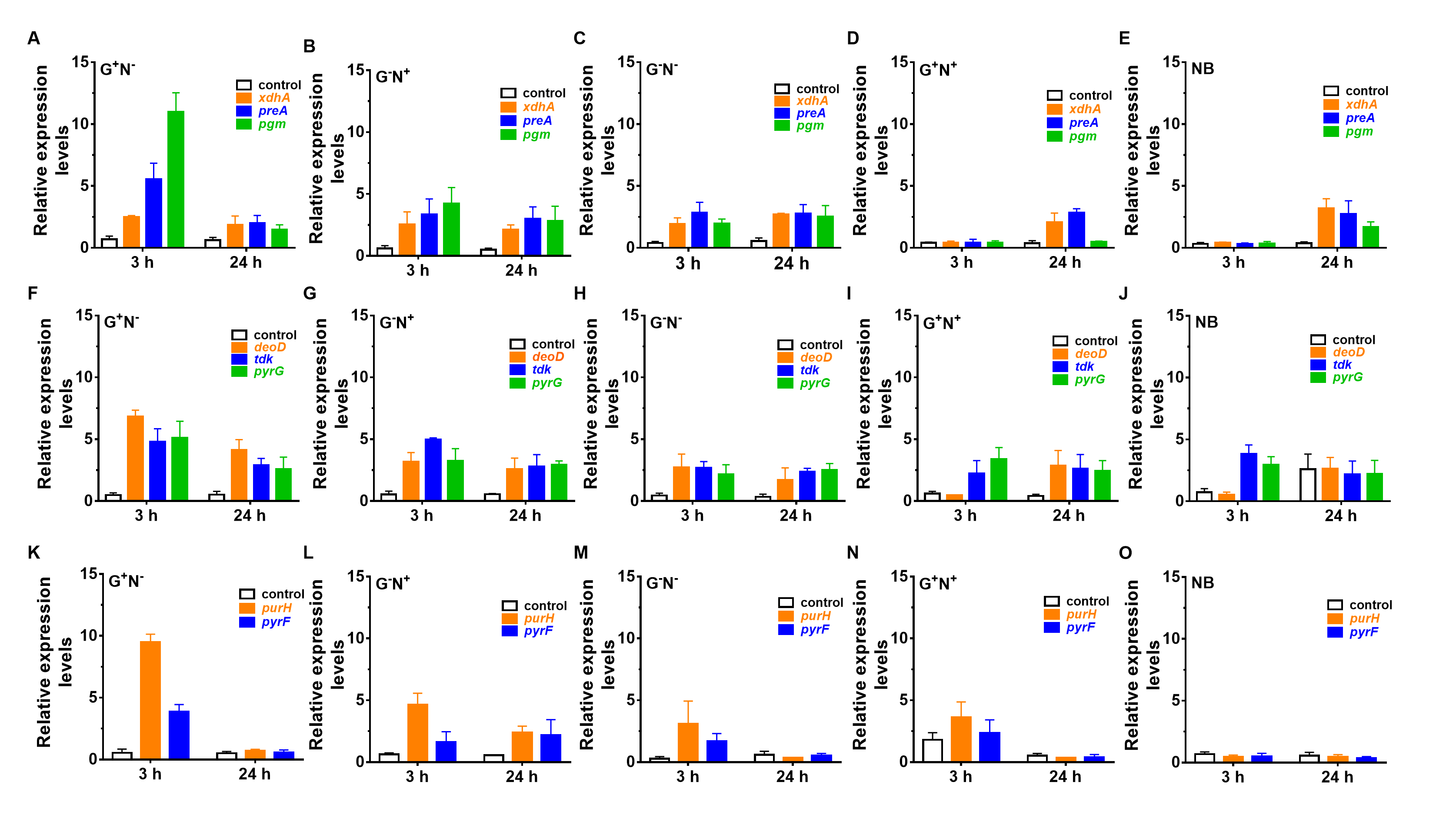

**Figure EV4. Gene expression of nuclotide decomposing and sythesis enzymes of *E. coli* after clutured in various M9 media containing DNA for 3 h and 24 h.**

(A) Gene expression of nuclotide decomposing enzymes *xdhA*, *preA*, *pgm* of *E. coli* after clutured in M9 medium lacking NH_4_Cl (G^+^Nˉ) containing DNA for 3 h and 24 h. The control is gene expression of purine de novo synthesis gene *xdhA*.

(B) Gene expression of nuclotide decomposing enzymes *xdhA*, *preA*, *pgm* of *E. coli* after clutured in M9 medium lacking glucose (GˉN^+^) containing DNA for 3 h and 24 h. The control is gene expression of purine de novo synthesis gene *xdhA*.

(C) Gene expression of nuclotide decomposing enzymes *xdhA*, *preA*, *pgm* of *E. coli* after clutured in M9 medium lacking glucose and NH_4_Cl (GˉNˉ) containing DNA for 3 h and 24 h. The control is gene expression of purine de novo synthesis gene *xdhA*.

(D) Gene expression of nuclotide decomposing enzymes *xdhA*, *preA*, *pgm* of *E. coli* after clutured in M9 medium (G^+^N^+^) containing DNA for 3 h and 24 h. The control is gene expression of purine de novo synthesis gene *xdhA*.

(E) Gene expression of nuclotide decomposing enzymes *xdhA*, *preA*, *pgm* of *E. coli* after clutured in M9 medium nutrient broth (NB) instead of M9 containing DNA for 3 h and 24 h. The control is gene expression of purine de novo synthesis gene *xdhA*.

(F) Gene expression of salvage nucleotide sythesis enzymes *deoD*, *tdk*, *pyrG* of *E. coli* after clutured in M9 medium lacking NH_4_Cl (G^+^Nˉ) containing DNA for 3 h and 24 h. The control is gene expression of purine de novo synthesis gene *deoD*.

(G) Gene expression of salvage nucleotide sythesis enzymes *deoD*, *tdk*, *pyrG* of *E. coli* after clutured in M9 medium lacking glucose (GˉN^+^) containing DNA for 3 h and 24 h. The control is gene expression of purine de novo synthesis gene *deoD*.

(H) Gene expression of salvage nucleotide sythesis enzymes *deoD*, *tdk*, *pyrG* of *E. coli* after clutured in M9 medium lacking glucose and NH_4_Cl (GˉNˉ) containing DNA for 3 h and 24 h. The control is gene expression of purine de novo synthesis gene *deoD*.

(I) Gene expression of salvage nucleotide sythesis enzymes *deoD*, *tdk*, *pyrG* of *E. coli* after clutured in M9 medium (G^+^N^+^) containing DNA for 3 h and 24 h. The control is gene expression of purine de novo synthesis gene *deoD*. Complete M9 medium.

(J) Gene expression of salvage nucleotide sythesis enzymes *deoD*, *tdk*, *pyrG* of *E. coli* after clutured in nutrient broth (NB) instead of M9 containing DNA for 3 h and 24 h. The control is gene expression of purine de novo synthesis gene *deoD*.

(K) Gene expression of *de novo* nucleotide synthesis enzymes *purH*, *pyrF* of *E. coli* after clutured in M9 medium lacking NH_4_Cl (G^+^Nˉ) containing DNA for 3 h and 24 h. The control is gene expression of purine de novo synthesis gene *xdhA*.

(L) Gene expression of *de novo* nucleotide synthesis enzymes *purH*, *pyrF* of *E. coli* after clutured in M9 medium lacking glucose (GˉN^+^) containing DNA for 3 h and 24 h. The control is gene expression of purine de novo synthesis gene *xdhA*.

(M) Gene expression of *de novo* nucleotide synthesis enzymes *purH*, *pyrF* of *E. coli* after clutured in M9 medium lacking glucose and NH_4_Cl (GˉNˉ) containing DNA for 3 h and 24 h. The control is gene expression of purine de novo synthesis gene *xdhA*.

(N) Gene expression of *de novo* nucleotide synthesis enzymes *purH*, *pyrF* of *E. coli* after clutured in M9 medium (G^+^N^+^) containing DNA for 3 h and 24 h. The control is gene expression of purine de novo synthesis gene *xdhA*.

(O) Gene expression of *de novo* nucleotide synthesis enzymes *purH*, *pyrF* of *E. coli* after clutured in nutrient broth (NB) instead of M9 containing DNA for 3 h and 24 h. The control is gene expression of purine de novo synthesis gene *xdhA*.

In Fig EV4, the expression of *xdhA* (purine decomposition gene), *preA* (pyrimidine decomposition gene) and *pgm* (deoxyribose decomposition gene) which are related to nucleotide decomposition was quantitatively analysed. When DNA was used to replace glucose or NH_4_Cl as carbon and nitrogen sources, these genes were all up-regulated (Fig EV4A-C). It indicates that in these media *E. coli* decomposes DNA and uses the produced small molecules as carbon and/or nitrogen source. In the media of M9 and nutrient broth, no nucleotide decomposition genes up-regulation was found after culturing for 3 h (Figure EV4D and E). While after 24 h, *xdhA* and *preA* related to nucleobase decomposition were up-regulated. The possible reason is that the grow speed of *E. coli* was not so quick in the first 3 h (see Fig 1 in the main text), and with the increase of culture time, DNA was ingested so that the nucleobases began to be decomposed and used as the nitrogen source. This result shows that DNA was utilized even in the presence of other nitrogen source, demonstrating that DNA is an excellent nitrogen source. On the other hand, *pgm* (deoxyribose decomposition gene) was not significantly up-regulated in M9 medium (Fig EV4D). The possible reason is that M9 medium contains enough glucose as a direct energy source, and the decomposition of deoxyribose was not stimulated to a high level.

The expression of *deoD* (deoxyribonucleoside synthesis gene), *tkd* (deoxynucleotide synthesis gene) and *pyrG* (CTP synthesis gene) which are related to nucleotide salvage synthesis was quantitatively analysed. These genes were up-regulated when DNA was used to replace glucose or NH_4_Cl as carbon and nitrogen sources (Fig EV4F-H). In the media lacking carbon and/or nitrogen sources, DNA has to be ingested and decomposed as the nutrient. At the same time, at the stages of either nucleotides or bases being produced, they can be directly used with the salvage pathway for synthesizing dNTPs and NTPs. In M9 and nutrient broth media, *deoD* related to deoxyribonucleoside synthesis was not significantly up-regulated (Fig EV4I and J). The possible reason is that there was not enough deoxyribose to synthesize deoxyribonucleosides. Strangely, genes related to nucleotide salvage synthesis in the control (no DNA) of nutrient broth were also up-regulated, which is hard to explain.

The expression of *purH* (purine *de novo* synthesis gene) and *pyrF* (pyrimidine *de novo* synthesis gene) which are related to nucleotide *de novo* synthesis was quantitatively analysed. These genes were all up-regulated after culturing for 3 h in various M9 media containing DNA (Fig EV4K-M). Especially in the medium lacking NH_4_Cl, there is a high level up-regulation of *purH* and *pyrF*, which indicates DNA was utilized as the material for nucleotide synthesis. DNA was decomposed and the produced small molecules were used as the nitrogen source to synthesize amino acids, which is the material for *de novo* RNA synthesis. It should be noted that deoxyribose cannot be oxidized to ribose, but ribose can be reduced to deoxyribose. Because only DNA was present, the *de novo* RNA synthesis pathway has to be active for growth. Interestingly, there was no significant expression in the nutrient broth. The possible reason is that nutrient broth may contain trace amounts of nucleotides, and *E. coli* can use them through the salvage synthesis pathway for growth.

**Table EV1.** The sequences of primers used in this study

| **Name** | **Sequence (5′→3′)** | **Length (nt)** |
| --- | --- | --- |
| *16S rRNA*-fw | CTGGAACTGAGACACGGTCC | 20 |
| *16S rRNA*-rev | GGTGCTTCTTCTGCGGGTAA | 20 |
| *endA*-fw | CAGTCGGTGAGGTGAATG | 18 |
| *endA*-rev | AGAGTGTCAGGTTGTATTGG | 20 |
| *comE*-fw | GGCTGGCAATGGTGATAG | 18 |
| *comE*-rev | GGCAACGATGATGAGGTAT | 19 |
| *hofQ*-fw | CACTACTGTTGATGCTGATAC | 21 |
| *hofQ*-rev | AGAGAATGTTGCCTTCCTG | 19 |
| *purH*-fw | GAACAGGAACTGCGTGAT | 18 |
| *purH* -rev | GCCAATGCCGATAGTCAT | 18 |
| *pyrF*-fw | TGTGTTCTGCTCAGGAAG | 18 |
| *pyrF*-rev | TCTACCGATTGCGTTACC | 18 |
| *deoD*-fw | GGAAGCGGCTGGTATCTA | 18 |
| *deoD*-rev | ATGTCGTTGAAGGTAGTCTG | 20 |
| *apt*-fw | GCAGTCTGACATCACCAT | 18 |
| *apt*-rev | GGCATGAAATCACGGAATT | 19 |
| *gpt*-fw | GACTGATGCCTTCTGAACA | 19 |
| *gpt*-rev | TGGTTGTCGTGATCGTAG | 18 |
| *hpt*-fw | GCGTCGGTGGAGATATTC | 18 |
| *hpt*-rev | GGAGTCAAGATTGCGGAAT | 19 |
| *upp*-fw | TAGCCTGCTGACTTACGA | 18 |
| *upp*-rev | ATACCGACAACGCTGATG | 18 |
| *xdhA*-fw | CCTTAGCCGTGAAGAGTG | 18 |
| *xdhA*-rev | GACAGAACATCCAGACTATAAC | 22 |
| *preA*-fw | TCGGCTGAAGGAAGATTAC | 19 |
| *preA*-rev | GCGGACAGGAGAAGTTAC | 18 |
| *pgm*-fw | GCTTCTATTGGCGGTCTG | 18 |
| *pgm-*rev | ATCTGCTTGCGATGTTCTT | 19 |
